# Structural connectome quantifies tumor invasion and predicts survival in glioblastoma patients

**DOI:** 10.1101/2021.03.09.434656

**Authors:** Yiran Wei, Chao Li, Zaixu Cui, Roxanne C. Mayrand, Jingjing Zou, Adrianna L.K.C. Wong, Rohitashwa Sinha, Tomasz Matys, Carola-Bibiane Schönlieb, Stephen John Price

## Abstract

Glioblastoma widely affects brain structure and function, and remodels neural connectivity. Characterizing the neural connectivity in glioblastoma may provide a tool to understand tumor invasion. Here, using a structural connectome approach based on diffusion MRI, we quantify the global and regional connectome disruptions in individual glioblastoma patients and investigate the prognostic value of connectome disruptions and topological properties. We show that the disruptions in the normal-appearing brain beyond the lesion could mediate the topological alteration of the connectome (*P* <0.001), associated with worse patient performance (*P* <0.001), cognitive function (*P* <0.001), and survival (overall survival: HR: 1.46, *P* = 0.049; progression-free survival: HR: 1.49, *P* = 0.019). Further, the preserved connectome in the normal-appearing brain demonstrates evidence of remodeling, where increased connectivity is associated with better overall survival (log-rank *P* = 0.005). Our approach reveals the glioblastoma invasion invisible on conventional MRI, promising to benefit patient stratification and precise treatment.

## Introduction

Glioblastoma is the most common primary malignant brain tumor in adults, characterized by diffuse infiltration into the surrounding tissue^1^. It is increasingly accepted that glioblastoma widely influences the brain structure and function beyond the focal lesion^2, 3, 4^. Microscopic evidence shows that glioblastoma can induce profound remodeling of neural connectivity^5^, while neuronal activity can promote tumor progression^6^. This bidirectional interaction underscores the promise of characterizing neural connectivity to better understand glioblastoma invasion, which may facilitate more accurate patient stratification for personalized management.

Diffusion MRI (dMRI) is a method to estimate the structural connectivity of the brain. It is more sensitive to detect occult tumor invasion, compared to the conventional T1-weighted and FLAIR images^7^. Evidence shows that dMRI can indicate the tissue signature of glioma^8^, offering values to evaluate invasiveness^9, 10^, detect peritumoral invasion^11, 12^, indicate subventricular zone involvement^13^, and predict molecular phenotypes^14^. These studies, however, have focused on the *focal* tumor, instead of the *systematic* disturbance of the brain.

The advance in neuroimaging has represented structural connectivity as a complex network, namely structural connectome^15, 16^. This approach models brain regions as nodes while the white matter connections among brain regions as edges. Graph theoretical analysis of the derived structural networks shows to characterize various neurological and psychiatric disorders^17, 18, 19^. Further, recent studies suggest that brain tumors may alter the connectome topology^20, 21, 22^. Initial evidence shows that the topological features derived from the structural connectome appear to outperform clinical parameters in survival prediction^23^. However, it requires further clinical validation whether the disruption of the structural connectome could be quantified for patient stratification. Of particular significance is whether the alteration of the neural connectivity could impact patient outcomes.

The purpose of this study was to characterize the disruption of the structural connectome in glioblastoma *systematically*. We hypothesized that glioblastoma could induce both focal and global disturbance to the structural connectome, leading to topological alteration of the brain and impact patient outcomes. We tested this hypothesis in two prospective glioblastoma cohorts. Firstly, we constructed the structural networks using the dMRI from glioblastoma patients and healthy controls. Secondly, we quantified the focal and distant disrupted connectome separately from each patient. Thirdly, we calculated the disruption indices and topological features and examined their significance in survival models. Lastly, we modeled the alteration of the preserved connectivity after removing the disrupted connectome, and investigated its significance on patient survival. The results reveal the widespread disruptions of the structural connectome, which lead to topological alterations and show prognostic value in glioblastoma patients.

## Results

### Subject characteristics

We included two patient cohorts in this study (**Supplementary Table S1,** see **Supplementary Methods** for a flowchart of patient inclusion). For the Discovery cohort, we recruited 136 patients for pre-operative MRI scanning. After excluding 19 patients according to the trial exclusion criteria, we included 117 of 136 (86.0 %) patients (mean age 59 years, range 22-75 years, 89 males) for analysis. Six patients (5.1 %) were lost to follow-up. The median overall survival (OS) was 392 (range 34-1932) days. The median progression-free survival (PFS) was 275 (range 13-1393) days.

For the Validation cohort, we initially recruited 49 patients. After excluding seven patients under the identical criteria with the Discovery cohort, we included 42 of 49 (85.7 %) patients (mean age 61 years, range 28-75 years, 34 males). The median OS was 335 (range 55-962) days. The median PFS was 246 (range 21-805) days. Two study cohorts showed no significant differences in clinical variables (**Supplementary Table S1**).

A cohort of 117 healthy age-matched subjects (mean age 59.9 years) was included as the control cohort. Another independent cohort of ten healthy controls (mean age 60.9 years) was included to generate a WM connection template. The above four cohorts showed no significant difference in age.

### Quantifying the strength of brain connectome

Tractography is a technique to measure the strength of the white matter (WM) connection by tracking the fiber pathway connecting different brain regions. However, directly performing tractography on the brain with a tumor may cause tracking failure or artifacts, e.g., an unrealistic belt of fibers surrounding the tumor^24^. To bypass the need to perform tractography on the lesioned brain, previous studies proposed to generate a template from healthy controls for tract localization, and the strength of WM connection was then robustly estimated by comparing patients to healthy controls^25, 26^.

We first derived a WM connection template among the 90 brain regions defined by the Automatic Anatomical Labeling (AAL) atlas^27^ (**Fig. 1A)**. As the strength of the WM connections can be measured by the fractional anisotropy (FA) value calculated from the dMRI, we generated an alignment-invariant tract representation through projecting FA voxels using tract-based spatial statistics (TBSS)^28, 29^. We further improved this approach by using the iterative projection under the guidance of tract orientation^30^. We also applied a histogram matching normalization method to reduce the variations across the subjects and imaging protocols^31, 32^. The skeletonized FA maps of patients and controls (**Fig. 1B)** were finally generated to produce the individualized strength matrices of WM connections (**Fig. 1C)**. We then calculated the strength vector of brain region disruptions by aggregating the WM connections.

**Fig 1.**
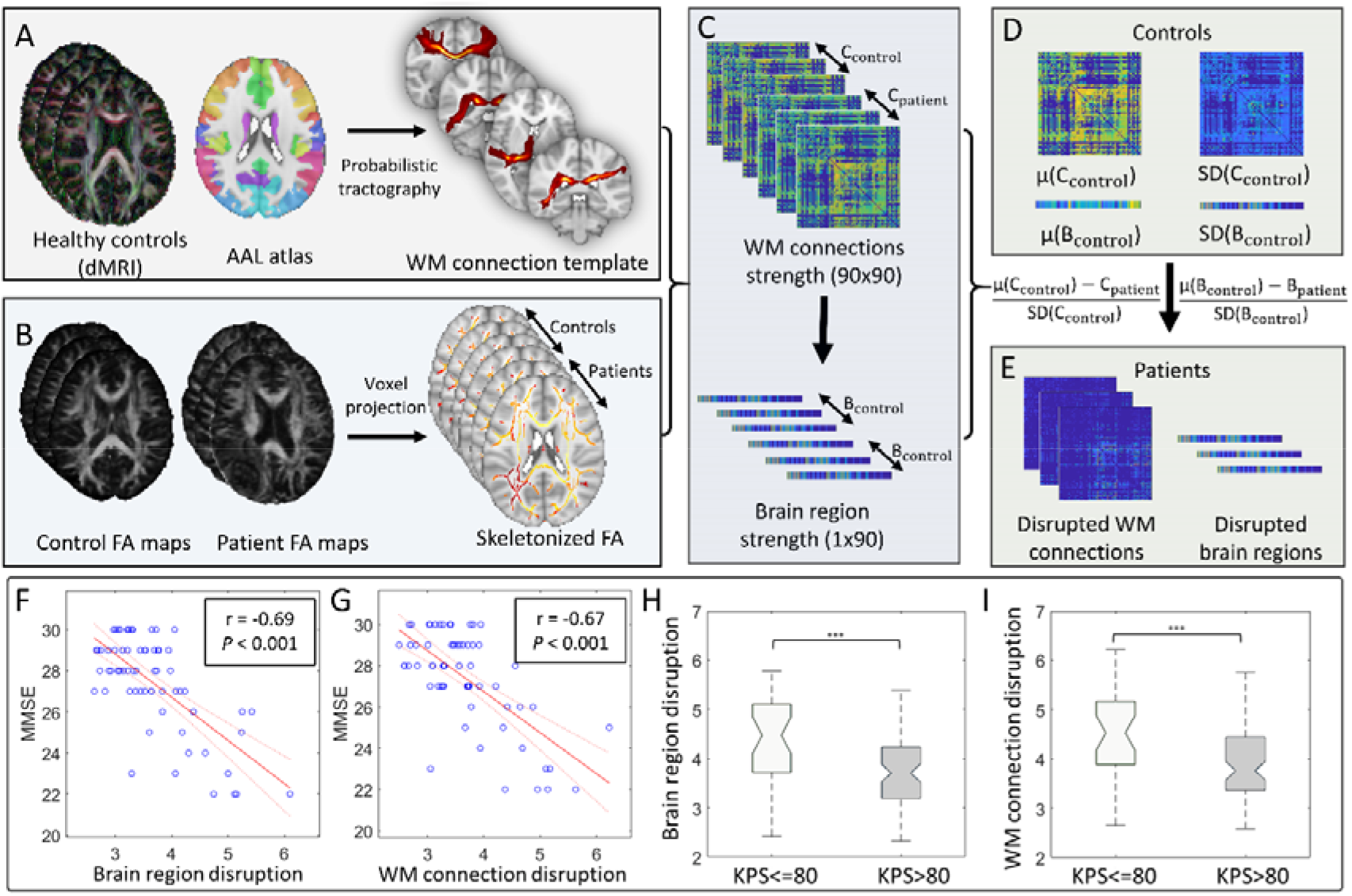
Quantifying connectome disruptions. **(A)** Probabilistic tractography is performed on the high-resolution dMRI of ten healthy controls to generate a template for the WM connections among the 90 regions on the AAL atlas. (**B)** Skeletonized FA maps are generated from both patients and age-matched healthy controls to estimate the WM connection strength, using an improved voxel projection procedure based on TBSS. The strengths of WM connections and brain regions are derived for healthy controls and patients **(C)**. By comparing the patients to the controls **(D)**, the disrupted connectome **(E)** in patients is identified as the WM connection (C_patient_) or brain region (B_patient_) with a strength that is 2SD (95% confidence) lower than the mean (μ(C_control_) or μ(B_control_)) of the control group. The disruption indices of WM connections and brain regions, calculated by averaging the disruption matrices/vectors, are both negatively correlated with the MMSE score **(F & G).** Higher disruption is associated with worse KPS (**H & I**). **AAL**: automated anatomical Labelling. **FA:** fractional anisotropy **WM:** white matter. TBSS: tract-based spatial statistics; **MMSE:** Mini-Mental State Examination**; KPS:** Karnofsky Performance Status. *P* value significant codes: *P* < 0.001 ***.

To rigorously assess the reproducibility of connectome strength estimation, we compared the strengths of a cerebellar tract (middle cerebellar peduncle, MCP) that is not affected by the supratentorial tumors in our cohorts. We found no significant difference in the strength of MCP across the patient and control cohorts (two-sample t-test and f-test, *P* > 0.05, **Supplementary Fig. S1**), suggesting that the connectome strength estimation was comparable across the cohorts.

### Global disrupted connectome demonstrates clinical significance

We calculated the disruption indices of both WM connections and brain regions as the standard deviation (SD)-normalized decrease in each patient, compared to the mean strength of the control group (**Fig. 1D**). We only considered the WM connection or brain region with a disruption index larger than 2SD (indicating 95% confidence) as significantly disrupted (**Fig. 1E**). We finally calculated the global disruption index in each patient by averaging the disruption of WM connections or brain regions, respectively.

To validate the clinical significance of the disruption indices, we tested their correlations with the Mini-Mental State Examination (MMSE) score. We observed that the global disruption of brain regions (r = −0.69, *P* < 0.001, **Fig. 1F**.) and WM connections (r = − 0.67, *P* < 0.001, **Fig. 1G**.) were both negatively correlated with the MMSE score. We further compared the patient subgroups stratified by the Karnofsky Performance Status (KPS) score of 80 as reported before^33^. We found that a worse KPS score was associated with higher disruptions of both brain regions and WM connections (both *P* < 0.001, **Fig. 1H**. **&** **I**). These results indicate that the global disruption indices were clinically robust, reflecting patient performance status.

### Identification of regional disrupted connectome

To address spatial tumor heterogeneity, we further calculated the regional disruption indices based on the global indices. We firstly segmented the tumors into contrast-enhancing regions (CE, the entire area within T1 contrast-enhancing rim) and peritumoral non-enhancing regions (NE, the hyper-intensities surrounding the CE on FLAIR images, **Fig. 2A**). Based on the segmented tumor regions, we categorized the disrupted WM connections as: **1) Direct** (crossing the tumor); **2) Indirect** (without crossing the tumor).

**Fig 2.**
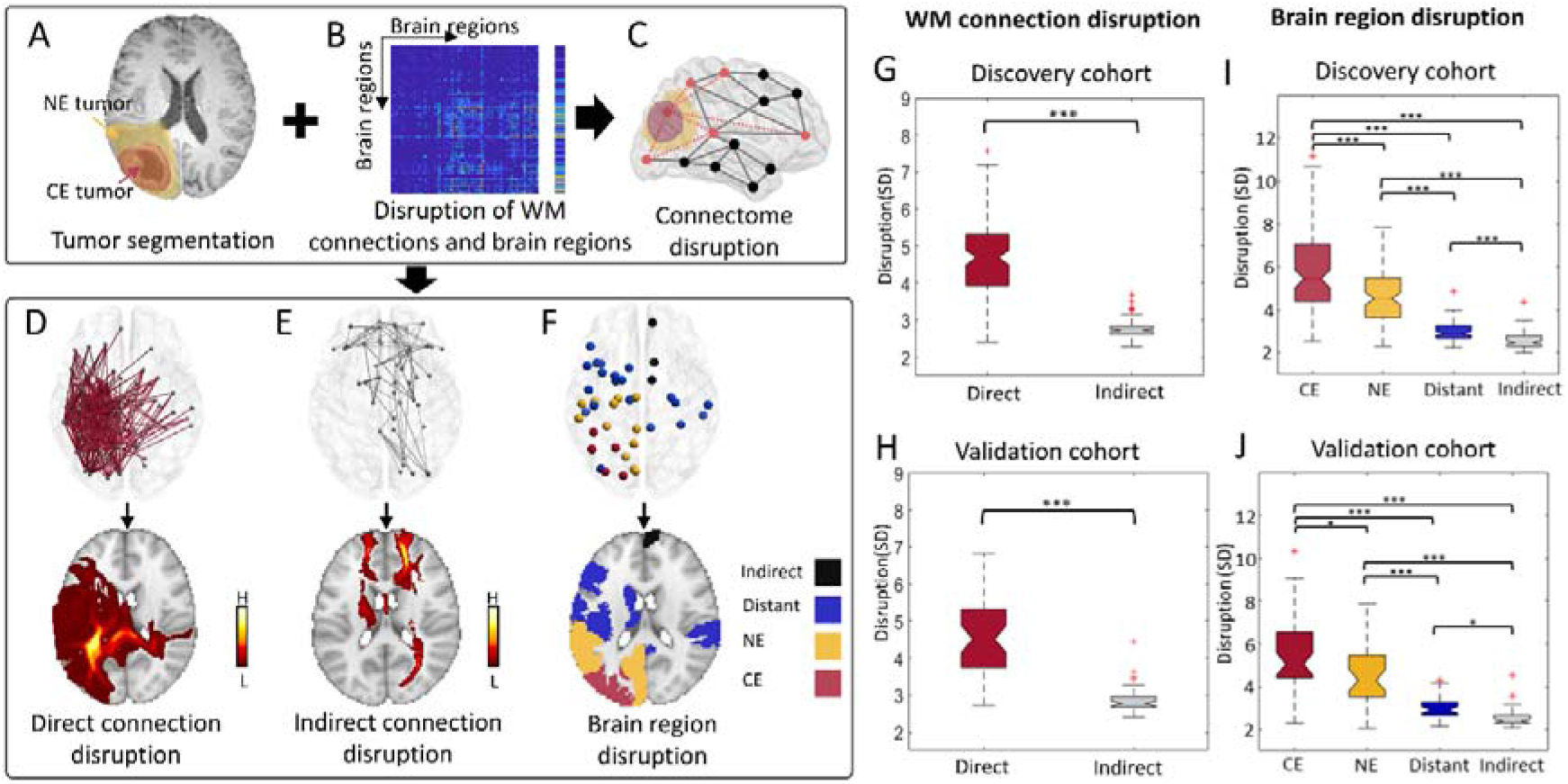
Quantifying regional disrupted connectome. The contrast-enhancing (CE, red) and non-enhancing (NE, yellow) tumors are segmented **(A)**. The disruption matrices of WM connections and disruption vectors of brain regions **(B)** are merged with the tumor segmentation to identify the regional disrupted connectome **(C)**. WM connection disruption is further classified as Direct **(D)** and Indirect **(E)** disruptions. The disrupted brain regions are classified into CE, NE, Distant, and Indirect disruptions **(F)**. In both Discovery and Validation cohorts, Direct connection disruption is higher than Indirect disruption **(G, H)**; Tumor regions (CE, NE) are more significantly disrupted than the normal-appearing brain (Distant, Indirect) **(I, J)**. **WM**: white matter. **MMSE:** Mini-Mental State Examination**; KPS:** Karnofsky Performance Status. *P* value significant codes: *P* < 0.001 ***, *P* < 0.01 **, *P* < 0.05 *.

The disrupted brain regions were similarly categorized as: **1) CE (** within CE tumor); **2) NE (** within NE tumor); **3) Distant (** within the normal-appearing brain beyond the tumor and connected to the tumor via WM connections**); 4) Indirect (** within the normal-appearing brain beyond the tumor without any WM connections with the tumor)

We defined the regional disruption index as the averaged disruption of each category (**Fig. 2D-F**). The two study cohorts showed no significant difference (**Supplementary Table S2**). In comparing disruptions of WM connections, we observed significantly higher Direct disruption (4.77 ± 1.56) than Indirect disruption (2.59 ± 0.39, *P* < 0.001, **Fig. 2G, Supplementary Table S3)**. Similarly for brain regions, focal tumor (CE: 5.83 ± 2.00; NE: 4.67 ± 1.33) were more significantly disrupted than the normal-appearing brain (Distant: 2.90 ± 0.71, Indirect: 2.56 ± 0.39, each *P* <0.001, **Fig. 2I**). The Validation cohort showed similar disruption patterns (**Fig. 2H** **&** **J**). These results correspond to our understanding of tumor invasion and support the robustness of the regional disruption indices.

### The regional disruptions are correlated with focal tumor volume

Pearson correlation tests showed that the WM connection disruptions in tumor (Direct) and normal-appearing brain (Indirect) were positively correlated (r = 0.44, *P* <0.001). Similarly, the disruption of Distant regions was positively correlated with that of tumor regions (Distant vs. CE: r = 0.43, *P* < 0.001; Distant vs. NE: r = 0.34, *P* = 0.028, **Supplementary Table S4**). Further, the tumor volume (measured by CE tumor) was positively correlated with the disruptions of both Direct connections (r = 0.52, *P* < 0.001) and Distant regions (r = 0.33, *P* < 0.001, **Supplementary Table S5**). Collectively, these data indicate that a larger focal tumor is associated with higher connectome disruption throughout the brain.

### The normal-appearing brain shows widespread disruption

We calculated the proportion of disrupted regions out of all brain regions. Noteworthy, in the patient group analysis, a higher proportion of Distant regions (16.8 ± 12.0%) was identified than the focal lesion (CE: 5.8 ± 5.1%, *P* < 0.001), recapitulated by the Validation cohort (**Supplementary Table S6**), supporting that the normal-appearing brain was widely disrupted.

Next, we explored the disruption probability of connectome in the patient population, calculated as the percentage of the patients with a specific region or connection disrupted. At the group level, the average disruption probability of Distant regions was higher (17.2 ± 9.0%) than focal lesion (CE: 11.8 ± 6.8%, *P* < 0.001), possibly due to the more extensive coverage of the Distant regions. This finding further confirmed that the disruption of brain regions was widespread beyond the lesion.

We further generated a tumor frequency map using the tumor segmentation of all the patients (**Fig. 3A**). The top five most likely disrupted Distant regions (**Fig. 3C)** were mainly in the low-frequency regions (See **Supplementary Table S8** for details). We also mapped the disrupted WM connections to the atlases of anatomical tract generated from 1,000 subjects in the UK Biobank^34^. Notably, the top five tracts most likely disrupted were mainly association tracts and close to the high-frequency regions (**Fig. 3B, Supplementary Table S7**), suggesting that the association tracts may mediate the tumor spread.

**Fig. 3.**
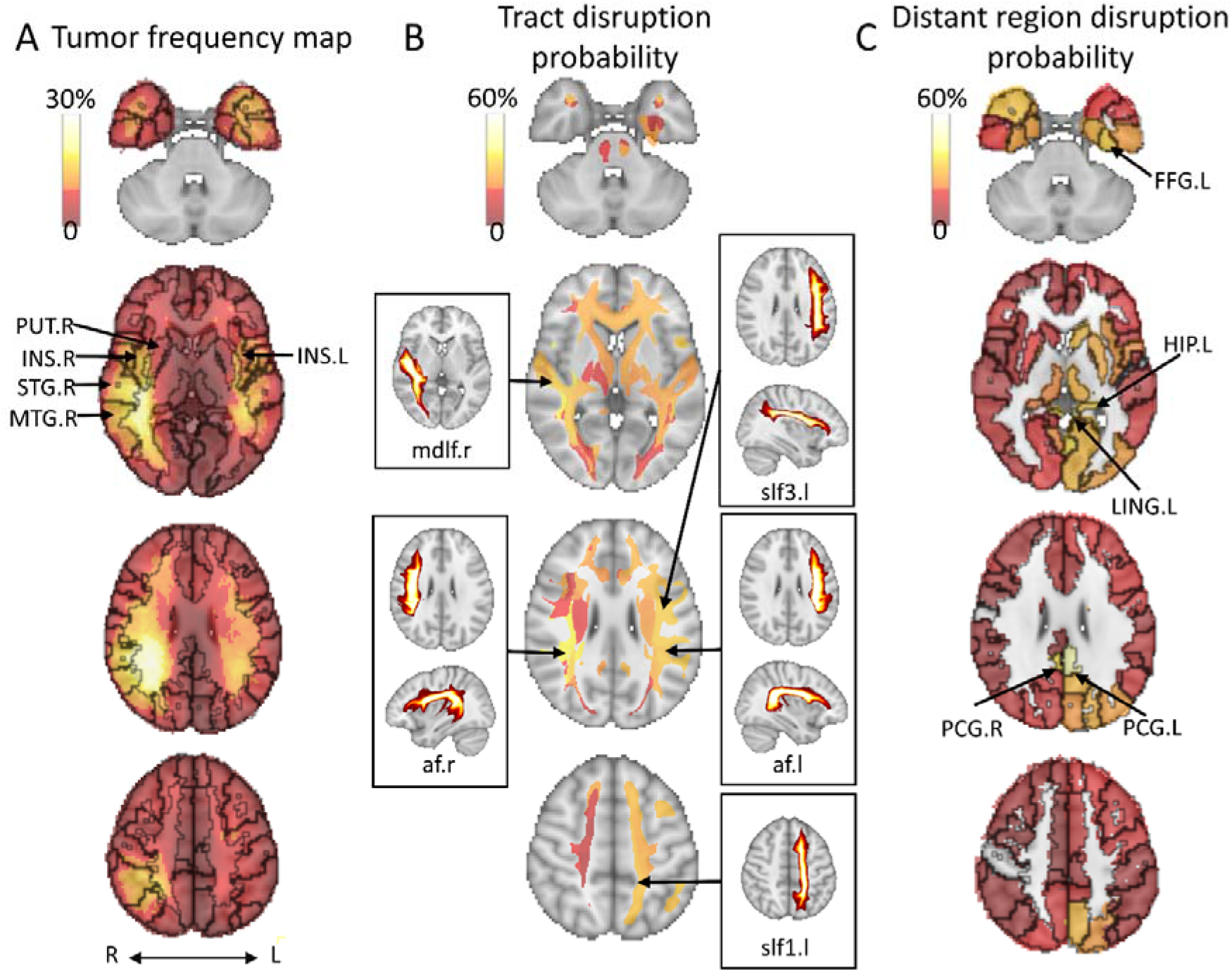
Tumor frequency map and disruption probabilities of anatomical structures. **(A)** Tumor frequency maps are generated using tumor segmentation. The top five disrupted focal brain regions include the right superior temporal gyrus (STG.R), right middle temporal gyrus (MTG.R), right insula (INS.R), left insula (INS.L), and right lenticular nucleus, putamen (PUT.R). **(B)** Top five disrupted anatomical tracts and their maximum intensity projection: right arcuate fasciculus (af.r), right middle longitudinal fasciculus (mdlf.r), left superior longitudinal fasciculus 3 (slf3.l), left arcuate fasciculus (af.l), left superior longitudinal fasciculus 1 (slf1.l). **(C)** The top five disrupted Distant brain regions include the left posterior cingulate gyrus (PCG.L), right posterior cingulate gyrus (PCG.R), left lingual gyrus (LING.L), left fusiform gyrus (FFG.L), and left hippocampus. L/l: left; R/r: right.

### The focal tumor alters the topological property of the connectome

To specifically investigate the topological alteration of the brain under tumor attack, we calculated the two most commonly used topological features: characteristic path length and clustering coefficient, which respectively reflect the efficiency of global communication and local information exchange in networks^35^. We observed that the characteristic path length of the patient networks was significantly higher than that of healthy controls (*P* < 0.001, **Fig. 4A**). In contrast, the clustering coefficient of patients was significantly lower than that of healthy controls (*P* < 0.001, **Fig. 4B**, **Supplementary Table S9**). These results reveal that tumor lesions could dramatically alter the topology property of the structural connectome.

**Fig. 4.**
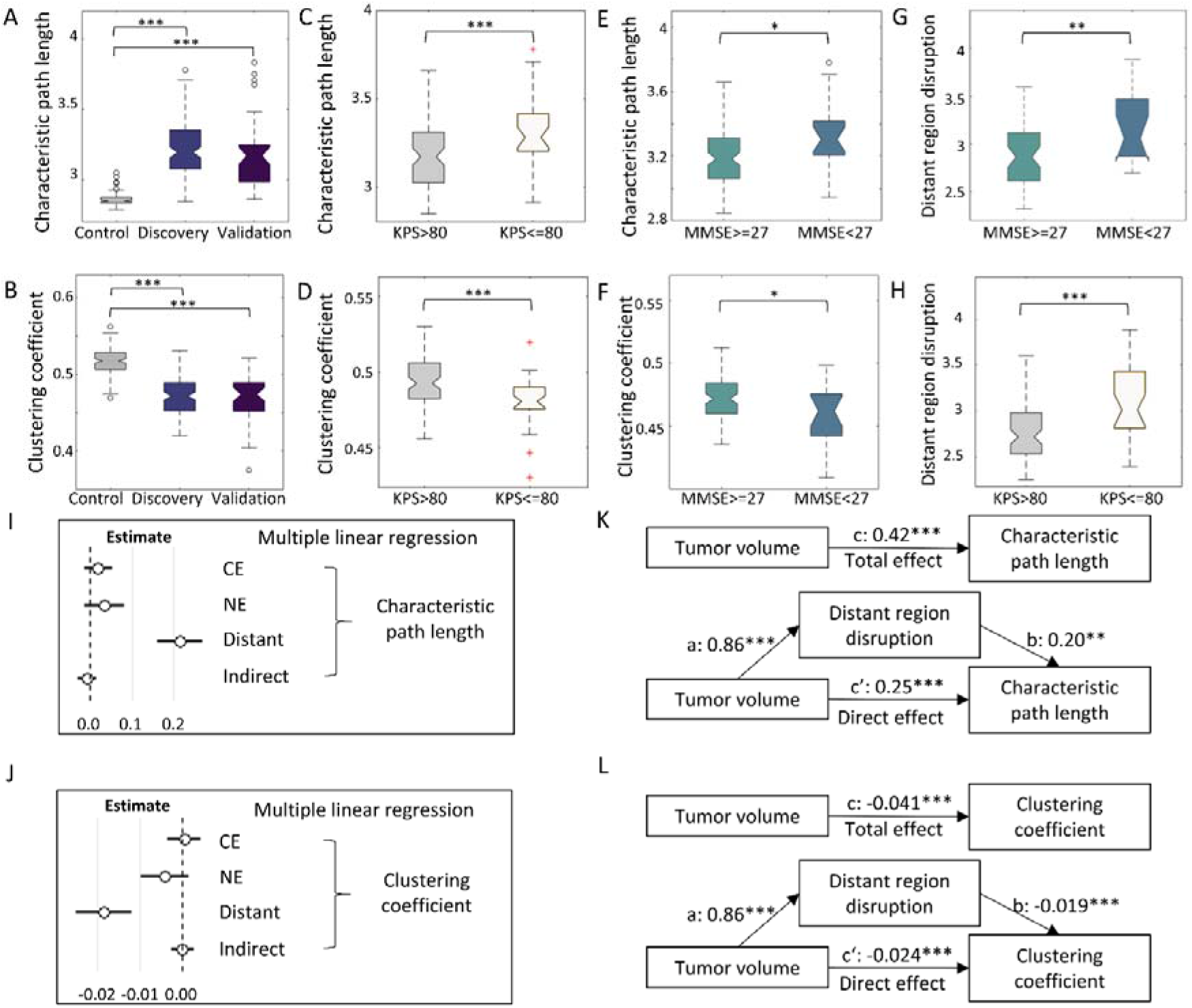
Topological alteration of the connectome. Patients show increased characteristic path length **(A)** and decreased clustering coefficient **(B)** than controls. Patient subgroups with worse pre-operative KPS **(C, D)** and MMSE **(E, F)** scores show increased characteristic path length and decreased clustering coefficient. Disruption of Distant regions is higher in the subgroups with worse MMSE **(G)** and KPS **(H)**, and it is the only significant predictor of characteristic path length **(I)** and clustering coefficient **(J)** in multiple linear regression. **(K)** The effects of tumor volume on characteristic path length are mediated by the disruption of Distant regions: total effect (c path) = 0.42, *P* < 0.001; direct effect (c’ path) = 0.25, *P* < 0.001; mediation effect (c – c’) = 0.17, *P* = 0.008. **(L)** The effects of tumor volume on clustering coefficient are mediated by the disruption of Distant regions: total effect (c path) = −0.041, *P* < 0.001; direct effect (c’ path) = −0.024, *P* < 0.001; mediation effect (c – c’) = −0.017, *P* < 0.001. **WM**: white matter, **MMSE** Mini-Mental State Examination**; KPS:** Karnofsky Performance Status. *P* value significant codes: *P* < 0.001 ***, *P* < 0.01 **, *P* < 0.05 *.

We next determined the clinical significance of connectome topology by comparing the topological properties of the patient subgroups stratified by MMSE and KPS scores. We found that the patients with lower MMSE or KPS scores presented lower clustering coefficient (MMSE: *P* = 0.012, KPS: *P* < 0.001, **Fig. 4C** **&** **D**) and higher characteristic path length (MMSE: *P* = 0.013, KPS: *P* < 0.001, **Fig. 4E** **&** **F**). Moreover, characteristic path length (r = 0.43, *P* < 0.001) was positively correlated with tumor volume, while clustering coefficient (r = −0.45, *P* < 0.001) was negatively correlated with tumor volume, indicating that a larger focal lesion may have a greater influence on the connectome topology (**Supplementary Table S5**).

### The disruption of Distant regions is associated with the topological alteration

To understand the relation between the regional disruption with topological properties, we performed a multiple linear regression, which revealed that the disruption of Distant regions was the only significant predictor of characteristic path length (Estimate = 0.21, *P* < 0.001) and clustering coefficient (Estimate = −0.018, *P* < 0.001, **Fig. 4I** **&** **J**). We then compared the disruptions of Distant regions in the patient subgroups stratified by the MMSE and KPS scores. We noticed that the patients with higher MMSE or higher KPS scores displayed significantly lower disruption of Distant regions (**Fig. 4G** **&** **H**), consistent with the distinct topological properties of these patient subgroups. The results imply the association between the disruption of Distant regions and connectome topology.

We further performed mediation analysis, which showed that tumor volume had both significant direct and indirect effects (mediated by the disruption of Distant regions) on characteristic path length (direct: *P* < 0.001, indirect *P* = 0.008) and clustering coefficient (direct & indirect *P* < 0.001) (**Fig. 4K** **&** **L**). The findings were confirmed by the Validation cohort (**Supplementary Fig. S3**).

### Topological features and disruption of Distant regions are prognostic

We evaluated the prognostic value of the disruption indices in log-rank tests. Stratified by the mean disruption of Distant regions (2.9), patients of higher disruption had worse survival than those of lower disruption (OS: median 293 vs. 449 days, *P* = 0.002, PFS: median 238 vs. 307 days, *P* = 0.019, **Fig. 5A**).

**Fig. 5.**
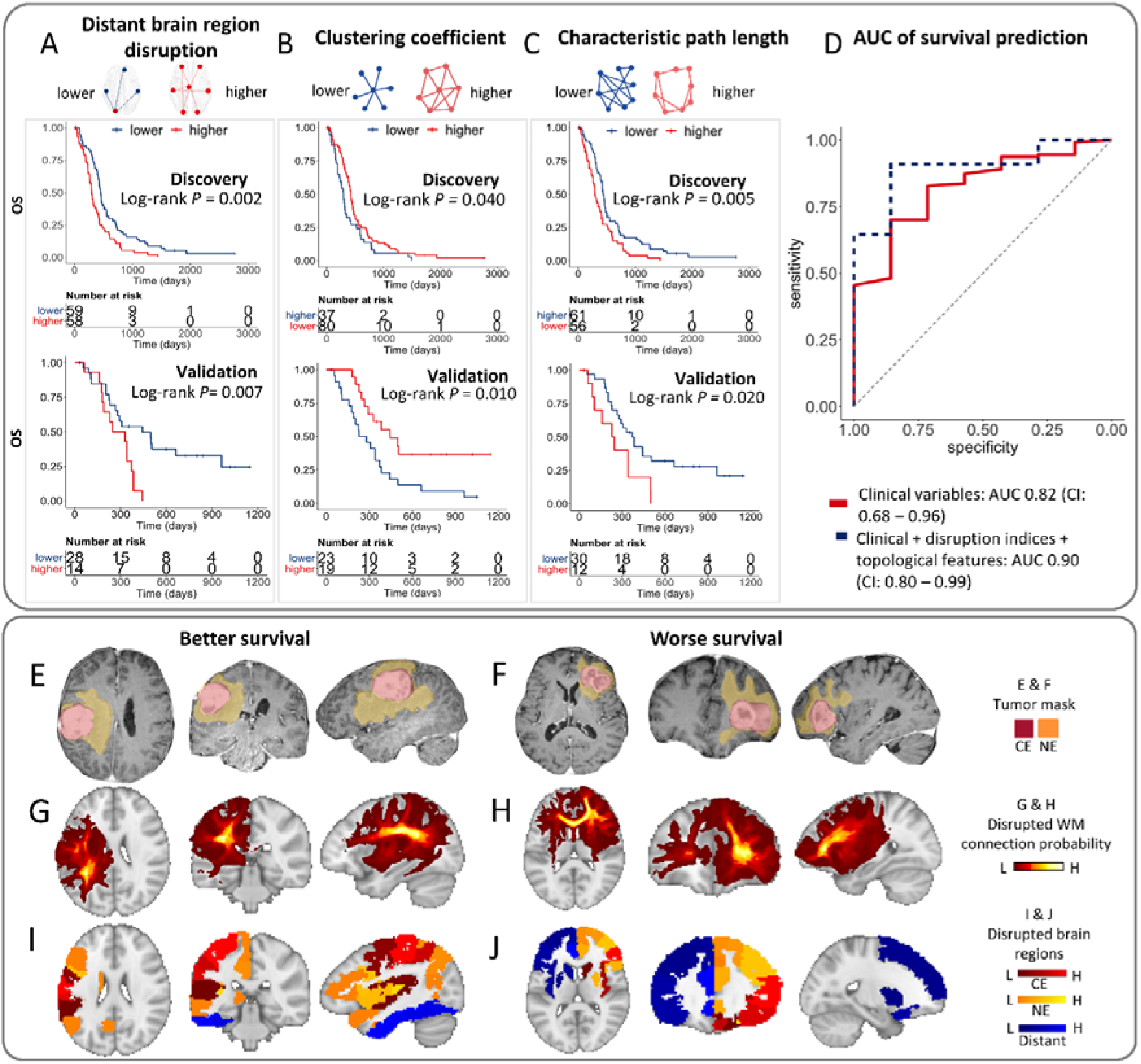
The prognostic value of disruption indices and topological features. **Top:** In both cohorts, higher disruption of Distant regions **(A)**, lower clustering coefficient **(B),** and higher characteristic path length **(C)** are associated with worse OS. **(D)** The model of predicting OS using clinical factors, disruption indices, and topological features shows improved AUC than clinical factors alone. **Bottom**: Two examples with better or worse survival (OS: 1555 vs. 317 days; PFS: 747 vs. 159 days). Both are IDH wildtype and MGMT unmethylated tumors of similar visible size in two males (aged 69 vs. 67 years). They both underwent complete resection followed by temozolomide chemoradiotherapy **(E & F).** Both patients have similar tumor sizes on post-contrast T1 (25.0 vs. 23.6 cm^3^). The patient with worse survival **(H)** has more widespread connection disruption beyond the visible lesion, compared to the patient with better survival **(G**); The disruption indices of Distant regions (blue) are 2.8 **(I)** and 3.0 **(J),** respectively. Their topological features are distinct (clustering coefficient 0.48 vs. 0.44: characteristic path length 3.17 vs. 3.31). **OS**: Overall survival. **HR**: Hazard ratio. **WM**: white matter.

Further, the subgroups stratified by the optimal cut-off of topological features (clustering coefficient: 0.46; characteristic path length: 3.20) had distinct survival. Precisely, the subgroup with a higher clustering coefficient had better survival than that with a lower clustering coefficient (OS: median 475 vs. 294 days, *P* = 0.040, PFS: median 306 vs. 238 days, *P* = 0.002, **Fig. 5B**). The subgroup with lower characteristic path length showed better survival than that with higher characteristic path length (OS: median 465 vs. 288 days, *P* = 0.005, PFS: median 312 vs. 244 days, *P* = 0.012, **Fig. 5C**). We further confirmed the findings in the Validation cohort using identical cut-offs.

We then evaluated the prognostic value of disruption indices and topological features in Cox models (**Table 1**). We observed that higher disruptions of Indirect connection (OS: HR = 1.36, *P* = 0.007; PFS: HR = 2.43, *P* = 0.046) and Distant regions (OS: HR = 1.46, *P* = 0.049; PFS: HR = 1.49, *P* = 0.019) were associated with worse survival. For topological features, higher clustering coefficient was associated with better survival (OS: HR = 0.63, *P* = 0.035; PFS: HR = 0.49, *P* = 0.002), while higher characteristic path length was associated with worse survival (OS: HR = 1.56, *P* = 0.035; PFS: HR = 1.82, *P* = 0.009, **Supplementary Table S10**).

**Table 1.** Univariate survival statistics of Discovery cohort.

| Feature | OS |  |  | PFS |  |  |
| --- | --- | --- | --- | --- | --- | --- |
| | HR | 95%CI | $P$ | HR | 95%CI | $P$ |
| **Clinical variables** |  |  |  |  |  |  |
| Age | 1.03 | 1.01-1.05 | **0.004** | 1.03 | 1.00-1.05 | **0.021** |
| Sex<sup>a</sup> | 0.81 | 0.52-1.25 | 0.333 | 0.75 | 0.47-1.19 | 0.217 |
| Performance<sup>b</sup> | 1.60 | 1.09-2.36 | **0.018** | 1.63 | 1.04-2.55 | **0.033** |
| IDH<sup>c</sup> | 0.59 | 0.25-1.36 | 0.211 | 0.52 | 0.22-1.21 | 0.131 |
| MGMT<sup>d</sup> | 0.77 | 0.52-1.14 | 0.196 | 0.68 | 0.43-1.07 | 0.094 |
| EOR<sup>e</sup> | 1.89 | 1.26-2.84 | **0.002** | 1.90 | 1.19-3.03 | **0.007** |
| Adjuvant treatment<sup>f</sup> | 0.21 | 0.13-0.34 | **<0.001** | 0.22 | 0.11-0.41 | **<0.001** |
| Tumor volume | 1.01 | 1.00-1.01 | **0.001** | 1.01 | 1.00-1.02 | **0.045** |
| Eloquent location<sup>g</sup> | 0.93 | 0.64-1.36 | 0.711 | 1.06 | 0.69-1.62 | 0.807 |
| Deep white matter<sup>h</sup> | 0.85 | 0.58-1.24 | 0.386 | 0.86 | 0.57-1.31 | 0.487 |
| **Disruption indices** |  |  |  |  |  |  |
| Direct connection | 1.07 | 0.87-1.30 | 0.790 | 1.08 | 0.88-1.32 | 0.464 |
| Indirect connection | 1.36 | 1.13-1.65 | **0.007** | 2.43 | 1.02-5.81 | **0.046** |
| CE regions | 0.95 | 0.86-1.04 | 0.667 | 0.96 | 0.86-1.06 | 0.418 |
| NE regions | 1.01 | 0.88-1.15 | 0.939 | 1.00 | 0.87-1.15 | 0.979 |
| Distant regions | 1.46 | 1.08-1.99 | **0.049** | 1.49 | 1.07-2.07 | **0.019** |
| Indirect regions | 1.06 | 0.87-1.29 | 0.790 | 0.80 | 0.83-1.28 | 0.795 |
| **Topological features** |  |  |  |  |  |  |
| Clustering coefficient | 0.63 | 0.42-0.93 | **0.035** | 0.49 | 0.30-0.78 | **0.002** |
| Characteristic path length | 1.56 | 1.06-2.29 | **0.035** | 1.82 | 1.16-2.84 | **0.009** |

(a). Female as the reference; (b). KPS 90-100 as the reference; (c). IDH wildtype as reference; (d) Unmethylated MGMT as reference; (e). Incomplete resection as reference; (f) Concurrent chemoradiotherapy (CCRT) as reference. (g). Non-eloquent location as reference. (h) Affected deep white matter as reference. **EOR**: extent of resection. **KPS**: Karnofsky Performance Score

We also evaluated the model performance in predicting OS using the disruption indices and topological features. The baseline model, including the above significant clinical variables (i.e., age, EOR, and adjuvant therapy), achieved an AUC of 0.82 (CI: 0.68-0.96). By adding the significant disruption indices and topological features into the baseline model, the AUC was improved at 0.90 (CI: 0.80 – 0.99, **Fig. 5D**). We presented two examples (**Fig. 5E-J)** with similar clinical variables but different disruption of Distant regions, topological features, and finally distinct survival (above and below the median, respectively).

In the multivariate model adjusting for all the significant clinical covariates from the univariate models, the disruption of Distant regions and topological features remained significant (**Fig. 6, Supplementary Table S10**). Their prognostic value was confirmed by the Validation cohort (**Supplementary Table S11 & S12**).

**Fig. 6.**
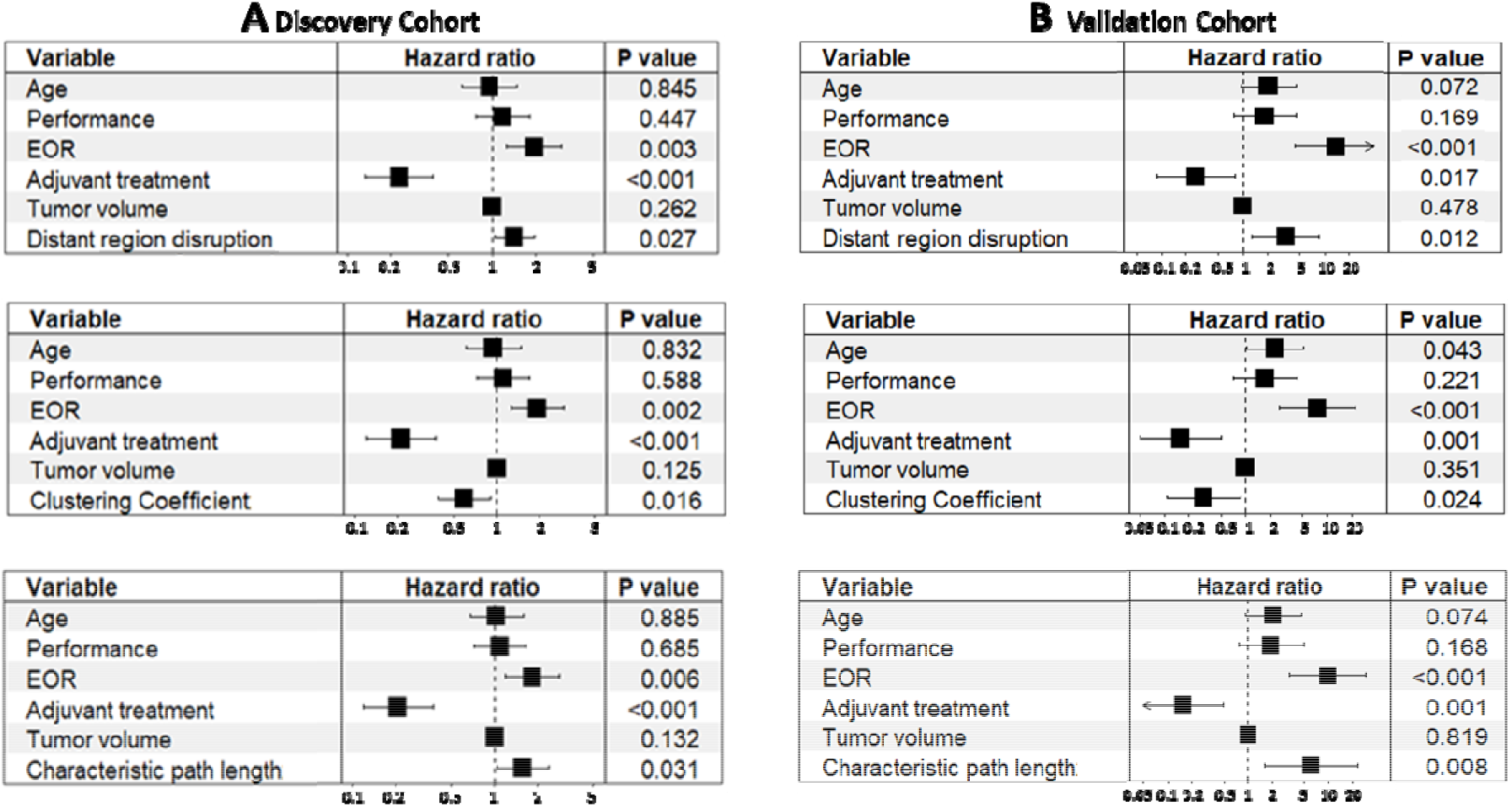
Forest plots of multivariate modeling of overall survival. For the Discovery **(A)** and Validation **(B)** cohorts, the higher disruption of Distant regions, higher characteristic path length, and lower clustering coefficient are associated with worse survival. Their prognostic value is independent of the significant clinical variables. **EOR**: extent of resection.

### The preserved connectome of Distant regions indicates patient survival

Given that the disruption of the Distant regions was associated with the topological alteration of the global connectome, we further investigated the preserved connectivity of the Distant regions, after excluding the significantly disrupted WM connections. Through pairwise comparison between patients and age-matched controls, we categorized the preserved connections into increased or decreased connectivity, respectively **(Fig. 7A)**. By aggregating the connections, we observed that 93.2% (109/117) patients displayed overall changes in connectivity, suggesting potential connectome remodeling. Among them, 24.7% (29/117) patients displayed overall increased connectivity, while 68.4% (80/117) patients showed overall decreased connectivity. We present two case examples with overall increased and decreased connectivity in the preserved connectome of the Distant regions, respectively **(Fig. 7A)**. The log-rank test showed that those patients with overall increased connectivity were associated with better survival (*P* = 0.005, **Fig. 7B**), confirmed by the validation cohort (**Supplementary Fig. 2**). The findings suggest that the remodeling towards a more integrated brain connectively, associated with the more robust connectome, may indicate better patient survival.

**Fig. 7.**
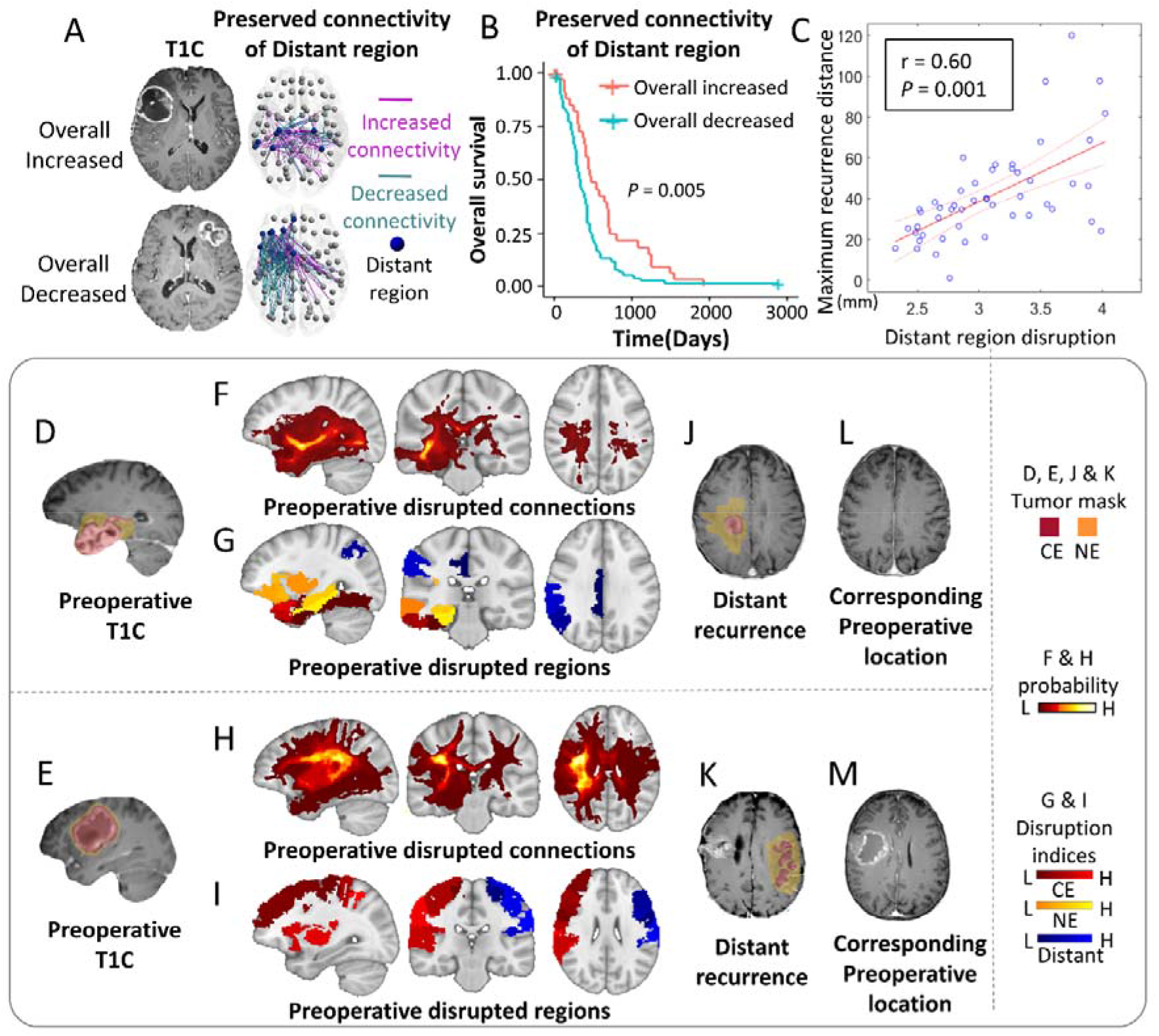
The disruptions of the Distant regions indicate patient survival and distant recurrence. **Top**: After removing the disrupted WM connections of the Distant regions, the preserved connections are categorized as increased or decreased connectivity in comparisons with healthy controls, and then aggregated to stratify patients. Two examples of overall increased and decreased connectivity are shown in **(A)**. The subgroup with overall increased connectivity shows better survival than overall decreased connectivity **(B)**. The disruption index of the Distant region is positively correlated with maximum recurrence distance **(C)**. **Bottom:** two examples of distant recurrence. Both patients present solitary visible lesions on pre-operative post-contrast T1 (T1C) images **(D, E)**, as well as widespread disrupted WM connections **(F, H)** and brain regions **(G, I**). In both patients, the distant recurrence location, either ipsilesional recurrence **(J)** or contralesional recurrence **(K),** corresponds to the Distant regions (blue), which are linked to the primary lesion via the WM connections shown in **(F, H**). A review of the pre-operative T1C images reveals no visible lesion in the recurrence location **(L, M)**.

### Disrupted connectome indicates tumor recurrence

Finally, we evaluated the usefulness of the disruption indices in indicating tumor recurrence after co-registering the follow-up recurrence scans to the pre-operative images. We found that the higher Distant region disruption was positively correlated with the furthest recurrence distance from tumor centroid (r= 0.60, *P* < 0.001, **Fig. 7C)**. We present two cases that showed distant recurrence in follow-up scans, where the disrupted Distant regions indicated occult tumor invasion invisible on the pre-operative MRI (**Fig. 7D-M**).

## Discussion

The present study employed a connectome approach to investigate the disruption of structural connectivity in glioblastoma. Our main findings include: 1) glioblastomas cause widespread disrupted neural connectivity beyond the focal lesion. 2) the disruption of the normal-appearing brain could mediate the alteration of connectome topology, associated with worse patient performance, and impact patient survival. 3) the preserved connectome demonstrates evidence of network remodeling that is associated with survival.

The finding that glioblastomas can cause widespread structural impairment is in line with the previous studies using resting-state fMRI, reporting that glioma induced widespread functional impairment.^3, 4^ The evidence supports that glioblastoma should be treated as a systematic disease rather than a local disease. Moreover, we found that only the disruption of Distant regions was associated with topological alteration and patient survival among all the regional disruptions, suggesting the importance of characterizing global neural connectivity.

In the anatomical mapping of the disrupted connectome, we found that the top disrupted Distant regions, e.g., posterior cingulate cortex and hippocampus, are essential structures of the limbic system, suggesting the propensity of the occult invasion affecting the limbic system. Moreover, the top affected anatomical tracts, e.g., arcuate fasciculus and superior longitudinal fasciculus, are long association tracts widely connecting separated gyri, suggesting that tumor invasion might spread through these tracts. Although at the macroscopic scale, our imaging findings may provide a perspective for previously reported neural-cancer interaction^36^.

We found that the connectivity measures could provide superior biomarkers for brain tumor stratification over conventional clinical factors, e.g., tumor location and volume. The network efficiency of the human brain generally reflects the integrity of brain function^37^. Glioblastoma patients displayed decreased network efficiency than healthy controls, likely due to tumor disturbance on brain function. Interestingly, our results suggest that the preserved connectome demonstrates evidence of remodeling. The increased connectivity, indicating a more integrated network and more robust function, is associated with favorable survival. Although the mechanism remains further explored, it could suggest the opportunities of understanding neural-cancer interaction for patient prognosis.

Our study has important clinical implications. Due to the remarkable heterogeneity of glioblastoma, the development of quantitative prognostic markers is crucial for precise diagnosis and treatment. The structural connectome and topological features confer a novel approach to investigate the systematic changes of neural connectivity in glioblastoma. It could enable us to understand the interaction between tumor invasion and neural connectivity, which promises to stratify patients more precisely and develop targeted therapeutics.

Our study has limitations. Firstly, the structural connectome can only directly measure the connectivity of connecting tracts. Although most brain regions are connected via tracts, certain functionally related regions may not be structurally connected. Future work could be improved by adding resting-state MRI and functional connectivity. Secondly, we only included primary glioblastoma who received first-line treatment in the trial setting. Therefore, molecular markers, i.e., IDH and MGMT methylation, were not significant as previously reported.

In conclusion, glioblastoma causes widespread impairment to the structural connectome. The invisible disruption on conventional MRI and connectome integrity are correlated with patient survival. Neural connectivity may provide a valuable tool for patient stratification and precise treatment.

## Methods

### Subjects

This study was approved by the local institutional review board. Informed written consent was obtained from all patients. Healthy control data were obtained from two open-source datasets, which have obtained ethical approval.

#### Glioblastoma patients

Patients with a radiological diagnosis of *de novo* supratentorial glioblastoma were prospectively recruited for surgical resection (Discovery: July 2010 – August 2015; Validation: July 2017 – October 2019) by the multidisciplinary team (MDT) central review. Patients were included in both cohorts following identical inclusion and exclusion criteria (see Supplementary methods). For both cohorts, patients were consecutively recruited, with data prospectively collected.

Patient pre-operative cognitive performance was tested using the Mini-Mental State Examination (MMSE) in the Discovery cohort. The MMSE score was dichotomized as <27 or >=27 as reported^38^. All glioblastoma patients underwent pre-operative 3D MPRAGE (pre-contrast [T1] and post-contrast [T1C]), T2-weighted FLAIR, and dMRI sequences.

#### Control cohorts

We included the age-matched subjects from the IXI datasets as healthy controls. The dMRI and T1 sequences of the cohort were available from https://brain-development.org/ixi-dataset/.

#### Template cohort

Healthy subjects of the template cohort were available from the Alzheimer’s disease Neuroimaging Initiative (ADNI, http://adni.loni.usc.edu/) for constructing an unbiased high spatial resolution template of WM connection. High angular resolution dMRI and T1 sequences were downloaded.

The scanning protocols of all the above subjects are detailed in the Supplementary Methods.

### Treatment

All patients underwent maximal safe surgery using 5-aminolevulinic acid fluorescence (5-ALA, Medac, Stirling, UK) and neuro-navigation (StealthStation, Medtronic, Fridley, MN, USA). According to the post-operative MRI within 72 hours, the extent of resection was assessed as complete or partial resection of enhancing tumor or biopsy. Adjuvant therapy was determined by the MDT, according to the standard treatment protocols based on the patient post-operative status. All patients were followed up after surgery according to the response assessment in neuro-oncology (RANO) criteria. Overall survival (OS) and progression-free survival (PFS) were used as endpoints.

### Tumor segmentation

All anatomical MRI, including T1, T2, and FLAIR, were co-registered to T1C images with an affine transformation, using the linear image registration tool (FLIRT) functions in the FMRIB Software Library (FSL)^39^. To segment the tumor, we applied a multi-scale 3D Deep Convolutional Neural Network^40^, implemented in the Cancer Imaging Phenomics Toolkit (CaPTk, https://cbica.github.io/CaPTk/index.html). A manual correction was performed using 3D slicer v4.6.2 (https://www.slicer.org/) by a neurosurgeon (XX) and a researcher (XX) after an initial training period and reviewed by an experienced neuroradiologist (XX). The final consensus was achieved to ensure inter-rater reliability.

### Connectome estimation

The complete pipeline of connectivity estimation includes three steps: 1) constructing group tract template, 2) producing individualized skeletonized FA map, 3) combining the WM connection template and FA skeleton to produce WM connection strength matrices.

#### WM connection template

An unbiased WM connection template in the standard space was generated by performing probabilistic tractography on the dMRI of the ten selected controls.

1. Cortical/subcortical regions of dMRI were parcellated into 90 brain regions according to AAL atlas 27 in the standard MNI-152 space^41^. Deformable registration was performed using the Advanced Normalization Tools (ANTs) ^42^. AAL atlas includes gray-white matter boundary to facilitate tractography.
2. Eddy currents and subject motions in dMRI were corrected using the FSL eddy tool (version 6.0.0). A crossing fiber model was then fitted to each control’s dMRI using the FSL function bedpostx. Probabilistic tractography between each pair of the 90 regions was subsequently performed using FSL Probtrackx2^43^. Each ROI was used as a seed (starting ROI) or target (ending ROI) once for tracking. For each pair of seed/target ROIs, 5000 streamlines were sampled from the seed mask. Only the streamlines that reached the target mask were retained. The tracking curvature threshold was set to 0.2 (80 degrees). Streamline samples were terminated when they have traveled 2000 steps with a step length of 0.5mm or entered the cortical/subcortical brain regions. Streamlines were discarded if they entered the cerebrospinal fluids (CSF) in the ventricle or re-entered the seed region.
3. For each healthy control, distribution maps were generated for all possible WM connections between the 90 cortical/subcortical brain regions. The distribution maps of the WM connections from all healthy controls were nonlinearly transformed to the MNI-152 standard space using ANTs and averaged to a mean WM connection distribution across controls using function fslmaths. The mean distribution was thresholded and binarized such that only the voxels with the top 5% probability in the WM connection were retained, providing a conservative pathway for the template.

#### Skeletonized FA map

To mitigate the partial-volume effect, we generated the skeletonized FA maps for estimating the strength of the WM connections in individual patients. The age-matched healthy subjects were selected as controls to reduce the bias from aging-related white matter pathology.

The dMRI was fitted with a tensor model to produce an FA map using the FSL diffusion toolbox (FDT)^44^. The FA maps were then nonlinearly co-registered to the MNI-152 space FA template using the deformable function of ANTs, which is shown to outperform the default deformable registration tools FNIRT ^45^ of TBSS in the co-registration of FA ^46^ and pathology-bearing T1 images^47^, and more importantly, could mitigate the deformation of the brain with tumor, by accounting for the tumor mass effect^48^. To minimize the bias of the signal-noise ratio introduced by the different MRI acquisition protocols, we normalized the FA map using the MRI intensity histogram-matching method^49^.

A standard space FA skeleton mask (FMRIB58 FA skeleton 1mm) was used as the target for FA voxel projection. The local maxima voxels from the FA map of patients and controls were projected to this skeleton mask using an improved TBSS projection guided by the tract orientation^30^. The generated individualized FA skeletons represent the center integrity of white matter tracts in subjects.

#### Constructing WM connection matrices

The WM connection matrix of each patient and control was estimated as the mean value of the tract segments in the individualized FA skeleton, constrained by the template of WM connection.

The columns and rows of each individualized WM connection matrix represent the brain regions in the AAL atlas, while the elements in the matrices (*C_ij_*) represent the strength of WM connection between the brain regions *i* and *j*. According to the graph theory, we calculated the strength *B_i_* for region *i*, by aggregating the connectivity strength of the WM connection *C_ij_* that are connected to brain regions *i* using the below formula:

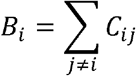

### Identification of significantly disrupted connectome

We first calculated the mean and SD of each connection strength across healthy controls. For individualized patient networks, we compared each connection strength to the mean connection strength of the control group. The significantly decreased connectome in patients was defined as a connection or brain region with the strength 2SD lower than the mean strength of the control group, where 2SD indicates 95% confidence.

### Identification of regional disrupted connectome

To address the intra-tumor heterogeneity, we categorized the disrupted WM connections and brain regions as below:

1. **Disrupted WM connections**

1. **Direct disrupted connections:** directly disrupted by tumor, traveling across the contrasting enhancing or non-enhancing tumor.
2. **Indirect disrupted connections:** disrupted without crossing the lesion region.
2. **Disrupted brain regions**

1. **Tumor disrupted regions:** the AAL brain regions within the tumor region and directly disrupted by the tumor. These regions were further categorized into the **CE** or **NE disrupted regions,** depending on the overlapping between tumor and AAL regions.
2. **Distant disrupted regions:** disrupted brain regions within the normal-appearing brain and connected to the tumor via WM connections.
3. **Indirect disrupted regions:** disrupted brain regions without any connections linked to the tumor.

We calculated each category of connection/region disruption as a disruption index by averaging the SD scaled decreases separately and generated five disruption indices for each patient.

### Tumor frequency map and disruption probability of anatomical structures

To use the segmented tumor masks to generate a focal tumor frequency map, we nonlinearly transformed all masks from individual patient space to the MNI-152 space using ANTs, with the voxel-wise tumor distribution density normalized at the group level.

To quantify the disruption probability of anatomical structures, we mapped the disrupted connectome to the prior atlases. For the anatomical tracts, we mapped the WM connections to the 42 anatomical tracts constructed from the 1,000 healthy subjects available from the XTRACT toolbox in FSL. The disruption probability for each brain region or tract was calculated as the percentage of patients with this brain region or tract disrupted.

### Topological features of the brain network

We calculated the clustering coefficient and characteristic path length using the Brain Connectivity Toolbox^16^. Briefly, the clustering coefficient measures the probability of two direct topological neighbors of a specific brain region being connected. The characteristic path length measures the average shortest path length of the network (see Supplementary Methods for detailed definition). To reduce the noise in feature calculation, we filtered the connectome with a population-consistency based strength threshold^50^.

### Statistical analysis

All analyses were performed in RStudio v3.2.3 (RStudio, Boston, USA) and MATLAB 2019b (The MathWorks Inc). The comparisons of disruption indices, topological features, and the performance subgroups were performed using a two-sample t-test. The correlation was tested using the Pearson correlation test. Multiple comparisons were adjusted by false discovery rate. Mediation analysis was performed using the R package ‘mediation’.

Survival analysis was performed using OS and PFS as the endpoints. Patients who were alive at the last known follow-up were censored. Disruption indices or topological features were dichotomized according to either median or the optimal cut-off value defined using the maximally selected rank statistics in the R package ‘Survminer’^51^, whichever was more significant. Kaplan-Meier survival curves were compared using the Log-rank test.

Cox proportional hazards regression accounted for all relevant clinical covariates, including O-6-methylguanine-DNA methyltransferase (MGMT) methylation status, isocitrate dehydrogenase-1 (IDH-1) mutation, sex, age, the extent of resection, adjuvant therapy, tumor volume. We also included two features from the VASARI feature set describing the involvement of eloquent cortex and deep white matter^52^ to account for the effects of tumor cortical/subcortical brain regions.

Receiver operating characteristic (ROC) curves were used to evaluate the accuracy of predicting OS, including the significant variables in the univariate models. To assess the prognostic values of tumor disruption and topological features, we fit a generalized linear model to calculate the region under the curve (AUC). The hypothesis of no effect was rejected at a two-sided level of 0.05.

## Acknowledgment

Part of the data were obtained from the Alzheimer’s Disease Neuroimaging Initiative (ADNI) database (adni.loni.usc.edu). As such, the investigators within the ADNI contributed to the design and implementation of ADNI and/or provided data but did not participate in the analysis or writing. The ADNI was launched in 2003 as a public-private partnership led by Principal Investigator Michael W. Weiner, MD. A complete listing of investigators can be found at http://adni.loni.usc.edu/. For up-to-date information, see www.adni-info.org.

Data collection and sharing were funded by the Alzheimer’s Disease Neuroimaging Initiative (ADNI) (National Institutes of Health Grant U01 AG024904) and DOD ADNI (Department of Defense award number W81XWH-12-2-0012). ADNI is funded by the National Institute on Aging, the National Institute of Biomedical Imaging and Bioengineering, and through generous contributions from the following: AbbVie, Alzheimer’s Association; Alzheimer’s Drug Discovery Foundation; Araclon Biotech; BioClinica, Inc.; Biogen; Bristol-Myers Squibb Company; CereSpir, Inc.; Cogstate; Eisai Inc.; Elan Pharmaceuticals, Inc.; Eli Lilly and Company; EuroImmun; F. Hoffmann-La Roche Ltd and its affiliated company Genentech, Inc.; Fujirebio; GE Healthcare; IXICO Ltd.; Janssen Alzheimer Immunotherapy Research & Development, LLC.; Johnson & Johnson Pharmaceutical Research & Development LLC.; Lumosity; Lundbeck; Merck & Co., Inc.; Meso Scale Diagnostics, LLC.; NeuroRx Research; Neurotrack Technologies; Novartis Pharmaceuticals Corporation; Pfizer Inc.; Piramal Imaging; Servier; Takeda Pharmaceutical Company; and Transition Therapeutics. The Canadian Institutes of Health Research is providing funds to support ADNI clinical sites in Canada. Private sector contributions are facilitated by the Foundation for the National Institutes of Health (www.fnih.org). The grantee organization is the Northern California Institute for Research and Education, and the study is coordinated by the Alzheimer’s Therapeutic Research Institute at the University of Southern California. ADNI data are disseminated by the Laboratory for Neuro Imaging at the University of Southern California.

